# Causal modulation of cortical amplitude coupling through dual-site amplitude-modulated tACS

**DOI:** 10.64898/2026.04.14.718451

**Authors:** Marina Fiene, Marcus Siems, Thorsten Kammerer, Till R. Schneider, Andreas K. Engel

## Abstract

**Background:** Intrinsic functional coupling at multiple temporal scales is a hallmark of human brain dynamics. Among these coupling modes, slow co-fluctuations of oscillatory amplitudes, termed amplitude coupling, are thought to represent a key organizing principle of the large-scale functional architecture, constraining and gating network activity. Yet, despite extensive correlational evidence, direct causal access to amplitude coupling remains limited, restricting insight into its functional relevance.

**Objectives:** Here, we investigated whether dual-site amplitude-modulated transcranial alternating current stimulation (AM-tACS) can selectively modulate interhemispheric amplitude coupling in human resting-state networks.

**Methods:** Twenty-eight participants received AM-tACS with a carrier frequency in the beta-band whose amplitude was modulated by low-frequency, scale-free dynamics. By applying dual-site AM-tACS either coherently or incoherently across bilateral parieto-occipital cortices, we tested whether stimulation could systematically enhance or disrupt amplitude co-fluctuations in the electrophysiological aftereffect.

**Results:** Incoherent AM-tACS significantly reduced interhemispheric amplitude coupling between targeted parieto-occipital cortices, with the strongest effects observed in the stimulated beta-band carrier frequency range. This modulation occurred independently of changes in local power or inter-areal phase coupling, indicating a selective effect of AM-tACS on amplitude-based connectivity. Moreover, reductions in amplitude coupling were correlated with the induced electric field strength, suggesting a dose-dependent relationship between stimulation intensity and coupling modulation.

**Conclusions:** Our findings demonstrate that dual-site AM-tACS can causally and selectively modulate amplitude coupling in the human brain. By establishing causal control over lasting amplitude coupling dynamics, this work provides a methodological foundation for future investigations into the functional and behavioral relevance of amplitude coupling in both healthy and pathological brain states.

**Highlights:**

- Dual-site AM-tACS selectively modulates amplitude coupling in humans
- AM-tACS was designed to mimic natural, scale-free amplitude fluctuations
- Stimulation effects are spatially confined to interactions between target regions
- E-field strength predicts the change in amplitude coupling, suggesting a dose-response relationship
- Amplitude coupling modulations are not mediated by band-limited power changes

## Introduction

Coupled slow co-fluctuations in the amplitude of neural oscillations, commonly referred to as amplitude coupling or amplitude envelope correlation, are a hallmark of large-scale functional networks in the human brain (Engel et al., 2013; Hipp et al., 2012; Siegel et al., 2012). These amplitude co-fluctuations are thought to orchestrate neural communication by shaping the temporal coordination of subnetworks relevant for cognition and behavior (Bruns et al., 2000; Engel et al., 2013; Siegel et al., 2012). Notably, amplitude coupling has emerged as a robust feature of spontaneous neural activity during resting state (Colclough et al., 2016; Hipp et al., 2012; Siems et al., 2016; Siems & Siegel, 2020). The amplitude correlation patterns derived from electrophysiological recordings closely correspond to resting-state networks identified using functional magnetic resonance imaging, underscoring a mechanistic link between electrophysiological dynamics and the hemodynamic brain network architecture (Brookes et al., 2011; Deco & Corbetta, 2011; Hipp et al., 2012; Hipp & Siegel, 2015; Mantini et al., 2007; O’Neill et al., 2015).

Given its central role in large-scale network coordination, alterations in amplitude coupling have been proposed as potential biomarkers of neural dysfunction and disease progression across multiple neurological and psychiatric disorders (Andreou et al., 2015; Maran et al., 2016; Raghavan et al., 2024; Siems et al., 2022). As demonstrated in magnetoencephalography resting-state recordings, diminished parieto-occipital alpha-band amplitude coupling has been observed in Parkinson’s disease, whereas Alzheimer’s disease was characterized by more widespread reductions in alpha- and beta-band amplitude coupling, (Boon et al., 2023; Schoonhoven et al., 2022; Stam et al., 2023; van Nifterick et al., 2024). These converging findings suggest disrupted amplitude coupling as a marker of impaired large-scale network coordination, with potential consequences for cognition and behavior.

Despite growing correlational evidence for a functional contribution of amplitude coupling to both healthy and pathological brain function, its causal role remains largely unknown. Addressing this question requires experimental tools capable of systematically manipulating coupling dynamics and assessing the resulting effects on neural activity. Transcranial alternating current stimulation (tACS) provides such an approach, enabling the frequency-specific modulation of neural oscillations through weak electric fields applied via scalp electrodes. Invasive recordings in animals have demonstrated that tACS can entrain neural activity, with neuronal spike timing aligning to the phase of the applied alternating current (Huang et al., 2021; Johnson et al., 2020; Krause et al., 2019, 2022; Vieira et al., 2020; Wischnewski et al., 2024). These findings were paralleled by evidence for neuronal entrainment effects in humans (Fiene et al., 2020, 2022; Neuling et al., 2012; Riecke et al., 2015; Schwab et al., 2019; Wischnewski et al., 2023), providing mechanistic evidence that tACS can reliably influence ongoing oscillatory dynamics in a controlled, frequency- and phase-specific manner.

A variant of this conventional sinusoidal tACS is amplitude modulated tACS (AM-tACS), in which a higher frequency carrier alternating current is modulated by a lower frequency envelope (Bächinger et al., 2017; Haslacher et al., 2024; Kasten et al., 2018; Minami & Amano, 2017; Thiele et al., 2021; Vieira et al., 2024; Witkowski et al., 2016). AM-tACS has mostly been applied using kilohertz-range carriers, assuming that physiological effects are primarily driven by the envelope frequency. Consistent with this view, Vieira et al. (2024) demonstrated that AM-tACS can entrain neuronal firing at the envelope frequency, albeit less effectively than conventional tACS. Although largely neglected, the carrier frequency may represent a key dimension for shaping AM-tACS effects. By jointly tuning carrier and envelope frequencies within biologically plausible ranges, AM-tACS could, in principle, engage neural dynamics across multiple temporal scales with greater specificity. Yet, it remains unresolved whether this approach improves the selective modulation of neural activity, and in particular, whether envelope phase manipulations during dual-site stimulation can causally influence large-scale amplitude coupling in electrophysiological recordings.

In the present study, we investigated whether dual-site AM-tACS can causally and selectively modulate amplitude coupling during resting state in humans. To this end, we tuned both the carrier frequency and the envelope dynamics to physiologically plausible parameters – at 17 Hz beta carrier frequency with scale-free amplitude modulation – and applied AM-tACS with either coherent or incoherent amplitude envelopes between bilateral parieto-occipital cortices. Using electroencephalography (EEG), we assessed the neurophysiological aftereffect of stimulation and hypothesized that amplitude coupling modulations would differ systematically between stimulation conditions, i.e., that incoherent AM-tACS would decrease interhemispheric amplitude coupling, whereas coherent AM-tACS would enhance it.

## Methods

### Participants

28 healthy participants (mean age 25.43 ± 4.38 years; 15 female; 13 male) were recruited from the University Medical Center Hamburg-Eppendorf, Germany. The sample size was chosen based on previous effect size observations in studies investigating neurophysiological effects of tACS on coupling metrics in humans (Alekseichuk et al., 2017; Helfrich et al., 2014; Polanía et al., 2012; Schwab et al., 2019). All participants were eligible for electrical brain stimulation, reported no history of psychiatric or neurological disorders and had normal or corrected-to-normal vision. Handedness was assessed via the short version of the Edinburgh Handedness Inventory (27 right-handed, 1 left-handed; mean laterality quotient 88.32 ± 36.38). The study was approved by the ethics committee of the Hamburg Medical Association (PV4908) and was conducted in accordance with the declaration of Helsinki. Written informed consent was given by all participants prior to participation and financial compensation was provided. One participant needed to be excluded due to technical issues related to the stimulation device. Therefore, the analysis includes data from 27 volunteers.

### Experimental design

Subjects participated in two experimental sessions (at least one day apart, 2.74 ± 2.57 days). Using a within-subject design, they received incoherent and coherent AM-tACS between bilateral parieto-occipital cortices in a counterbalanced, randomized order, with participants being blind towards the AM-tACS stimulation sequence. Participants were seated in a dimly lit, electrically shielded and sound-attenuated EEG chamber, facing a grey screen. They were instructed to fixate on a central fixation cross displayed on the screen. Each session started with a 10 min block of resting-state EEG recording without electrical stimulation, followed by four blocks of 10 min AM-tACS, each directly followed by 5 min (in the last block 10 min) resting-state EEG recording (Fig. 1A). Thereby, we captured the outlasting aftereffect of electrical stimulation on neurophysiological brain activity. Between the blocks, participants had a short break of about 8 min.

**Fig. 1.**
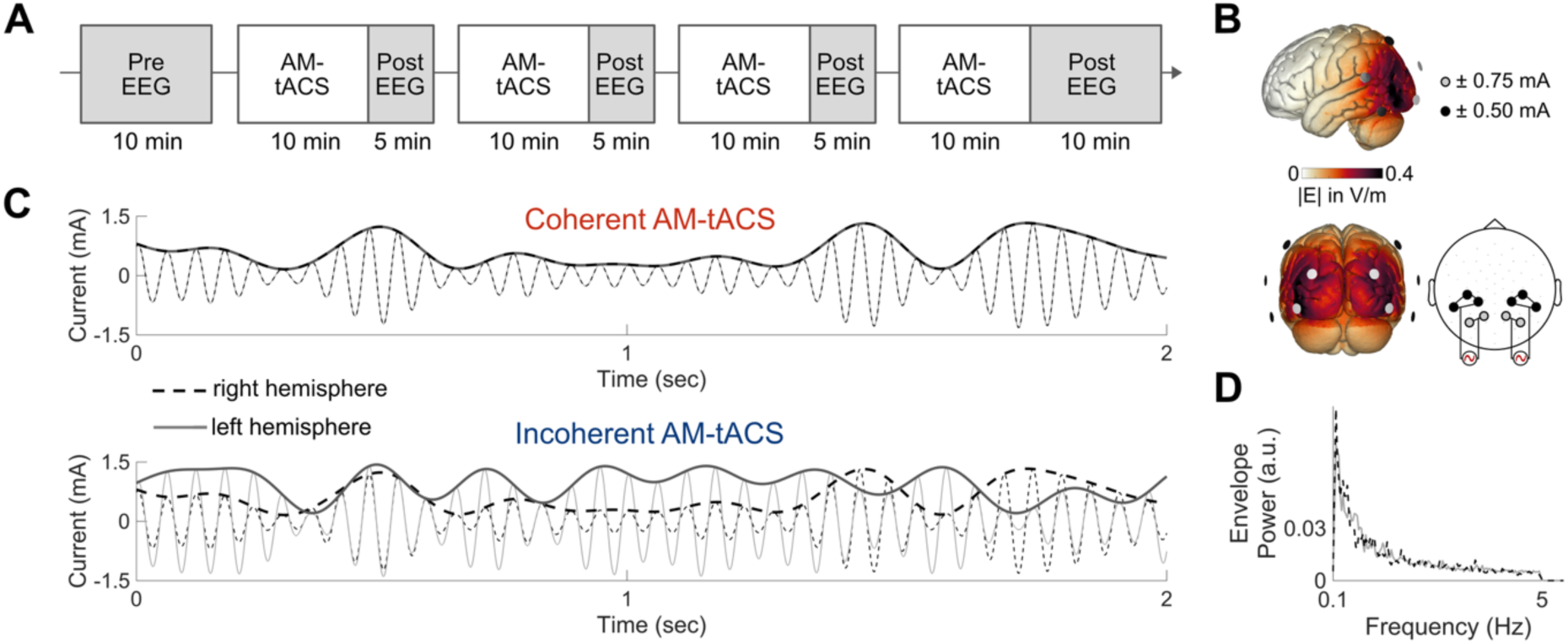
Experimental setup. (A) Timeline of resting-state EEG recordings. The pre resting-state EEG recording at baseline was followed by four blocks of AM-tACS, each directly followed by post resting-state EEG recordings. (B) Dual-site AM-tACS was applied at 3 mA peak-to-peak using two 2 x 3 multi-electrode montages over the left and right parieto-occipital cortices. Estimated maximum electric field strength was about 0.4 V/m in the AM-tACS target region. (C) Stimulation current waveforms of the two AM-tACS conditions applied to the left hemisphere (gray) and to the right hemisphere (black dashed), plotted for 2 sec. The amplitudes of the 17 Hz carrier signals were either modulated in a coherent or incoherent manner between hemispheres. (D) Spectrum of the AM-tACS amplitude envelopes, showing scale-free dynamics between 0.1 – 5 Hz.

To ensure that participants stayed awake during the course of the session, we included a simple vigilance task applied only during AM-tACS. Two tones of high or low pitch were played in randomized order every 5 – 10 sec. Participants’ task was to indicate the pitch of the tone by pressing stress balls with their left or right hand. The assignment of the left and right hand to the tone pitches was counterbalanced both within and across participants.

In the end of each session, peripheral AM-tACS side effects were systematically quantified via a questionnaire assessing the perceived strength of tactile sensations during electrical stimulation (itching, warmth, stinging, pulsing, pain) on a five-point scale as “absent”, “weak”, “moderate”, “pronounced”, or “intense”, as well as the strength of cognitive fatigue as “very awake”, “awake”, “moderate”, “tired”, or “very tired” over the course of the testing session.

### Amplitude-modulated transcranial alternating current stimulation

Multi-electrode AM-tACS was applied via 10 Ag/AgCl electrodes with 12 mm diameter (NG Pistim, Neuroelectrics) using the Starstim20 stimulator (Neuroelectrics, Barcelona, Spain). Electrodes were positioned over bilateral parieto-occipital cortices (Fig. 1B), regions previously shown to exhibit pronounced interhemispheric amplitude coupling during resting state in the human brain (Hipp et al., 2012; Siems et al., 2016). Electric field strength and distribution were estimated using SimNIBS 4.1.0 (Thielscher et al., 2015). To minimize transcutaneous sensations during electrical stimulation, EMLA cream (2.5 % lidocaine, 2.5 % prilocaine, Aspen, Germany) was applied for local anesthesia one hour prior to the start of the experiment. Stimulation electrodes were then prepared using Signa gel (Parker Laboratories, USA), and impedance was kept below 20 kΩ. Stimulation intensity was set to 3 mA peak-to-peak, with 7 sec ramp-up and 3 sec ramp-down periods, for a total duration of 40 min per session, divided into four blocks.

The AM-tACS waveform consisted of a 17 Hz carrier, amplitude-modulated by a low-frequency envelope spanning 0.1 – 5 Hz (Fig. 1C and D). Two envelope signals, one for stimulation of each hemisphere, were generated from filtered pink noise following these steps: 10 min pink noise data segments were bandpass filtered (0.1 – 5 Hz, fir1), normalized by the signal’s Hilbert transform magnitude, filtered again (0.1 – 5 Hz, fir1), and rescaled between 0 and 1 by subtracting the minimum and dividing by the maximum value. This procedure produced envelopes with a scale-free power spectrum and a tapered uniform amplitude distribution. The mean magnitude squared coherence between the two generated envelopes in the frequency range from 0.1 – 5 Hz was 0.028. These two envelopes were constructed once and applied consistently across all participants for coherent and incoherent stimulation.

### Electrophysiological recording and data preprocessing

EEG was recorded from 64 Ag/AgCl electrodes mounted in an elastic cap (EasyCap, Germany). The EEG was referenced to the nose tip and the electrooculogram was recorded by two electrodes placed below the eyes. Data acquisition was performed using the BrainAmp DC amplifiers (Brain Products GmbH, Germany) and the corresponding software (BrainVision Recorder 1.20). Electrodes were prepared using abrasive conducting gel (Abralyt 2000, EasyCap, Germany) keeping impedances below 20 kΩ. Data were recorded with an online passband of 0.016-250 Hz and digitized with a sampling rate of 1000 Hz.

EEG data were preprocessed in MATLAB using the analysis toolbox EEGLAB (Delorme & Makeig, 2004), FieldTrip (Oostenveld et al., 2011), as well as custom-made scripts. The technical offset artifact of the Starstim stimulator occurred at 117.13 ± 58.54 ms after the end of electrical stimulation, thus, data were trimmed accordingly to exclude this artifact from the analysis time window. Data were high-pass filtered at 0.5 Hz, lowpass-filtered at 48 Hz and down sampled to 500 Hz. Channel epochs that remained clipped shortly after AM-tACS offset due to amplifier saturation were interpolated by all other non-clipped channels using spherical interpolation (mean number of clipped channels 2.54 ± 1.73). To automatically identify electrodes with high noise levels, channels whose standard deviation exceeded three times the median standard deviation across all channels per participant were excluded and replaced by spherical interpolation based on the remaining channels (mean number of noisy channels 1.56 ± 1.72). Any residual non-stereotyped artifacts were then marked based on visual inspection of the data.

Independent component analysis (ICA) was computed on the artifact cleaned dataset using the infomax ICA algorithm (Bell & Sejnowski, 1995). ICA weights were then applied to the continuous dataset (including artifactual epochs) and artifactual components related to eye-movements, cardiac, and muscle activity were identified based on visual inspection of the components’ time course, spectrum and topography (Chaumon et al., 2015; Debener et al., 2010). On average, 25.56 ± 4.77 components were excluded.

### Source projection

We used a boundary element method volume conduction model, which we computed with FieldTrip based on the template structural NMRI225 provided by Kreilkamp et al. (2023) and computed the forward model for 457 equally spaced cortical sources (about 1.2 cm distance, at 0.7 cm depth below the pial surface) (Hipp & Siegel, 2015). Sensor level EEG data were projected into source space using dynamic imaging of coherent sources (DICS) linear beamforming (Gross et al., 2001). This method reconstructs activity at locations of interest with unit gain while maximally suppressing contributions from spatially distinct sources.

### Spectral analysis

Spectral estimates of the time-domain EEG signal were computed using Morlet’s wavelets (Goupillaud et al., 1984)

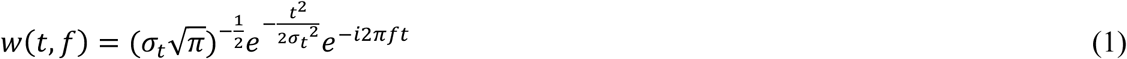

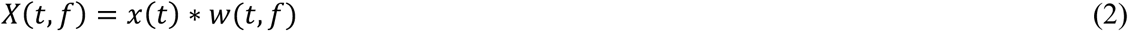

where *f* is the carrier frequency, *σ_t_* is the temporal standard deviation. The time-frequency estimate *X*(*t*, *f*) of a signal *x*(*t*) was then computed by convolution with the wavelet kernel *w*(*t*, *f*). The frequencies were linearly spaced between 1 to 30 Hz and the bandwidth of wavelets was set to 0.5 octaves. We computed spectral estimates on the continuous EEG data epochs to ensure robust amplitude estimation.

### Coupling analysis

As a measure of interhemispheric amplitude coupling, we computed power envelope correlations between orthogonalized neural signals (Hipp et al., 2012) on sensor and source level. Orthogonalization is necessary to account for volume conduction and, thus, overestimation of short-distance correlations. We first orthogonalized two complex signals at each time point by

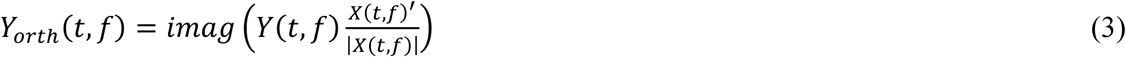

*X* and *Y* are two signals described by a function of time and frequency. *X*′ is the complex conjugate of *X*. The *imag* operator describes the imaginary part of the signal. Second, we computed the pairwise Pearson’s correlations of the log-transformed power envelopes of the signals *X* and *Y_orth_* for artifact free data segments only (artifactual data epochs ± half a wavelet length were removed). As orthogonalization between two time-series can be done in two directions (*X* to *Y* and *Y* and *X*), we computed power envelope correlations for both directions and averaged the two values per electrode pair.

To assess phase coupling, for each electrode pair the absolute value of the imaginary part of coherency (Nolte et al., 2004), i.e., imaginary coherence, was computed as

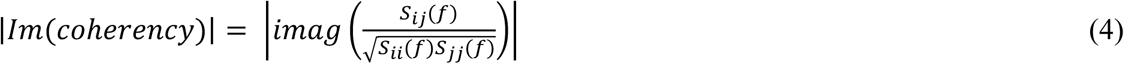

Here, the auto- and cross-spectra *S_ij_*(*f*) = 〈*x_i_*(*f*)*x_j_*′(*f*)〉 of electrodes *i* and *j* were computed from the complex signals derived from Morlet’s wavelet convolution. As for amplitude coupling, imaginary coherence was computed on artifact-free data only.

### Statistical analysis

According to our main hypothesis, we assessed the effect of dual-site AM-tACS on the change in interhemispheric amplitude coupling. To accurately assess the spatial structure of correlation values while accounting for interindividual variability in overall coupling strength, we standardized (z-scored) coupling values per subject and carrier frequency. For between-condition comparisons, z-scoring was performed across all channels, epochs and both conditions per subject and frequency, whereas for within-condition comparisons, z-scoring was conducted across channels and epochs, but separately within each condition, subject and frequency. As we expected to observe stimulation effects specifically in the brain area targeted by AM-tACS, we defined a parieto-occipital electrode cluster surrounding AM-tACS electrodes. For homologous electrode pairs within this cluster, we computed the difference scores (post minus pre AM-tACS) in amplitude coupling and applied cluster permutation statistics to detect carrier frequency clusters depicting significant differences in amplitude coupling within or between AM-tACS conditions. A cluster was defined as the sum of *t*-values of neighboring carrier frequencies showing a significant change in amplitude coupling (*p* < .05). The permutation distribution for within condition comparison was generated by randomly shuffling pre and post AM-tACS amplitude values within participants, consistently for all epochs and channels per participant, across 10.000 iterations. For between condition permutation, the assignment of incoherent or coherent condition was randomly shuffled per participant, again consistently for all epochs and channels per subject. The observed cluster was considered statistically significant when the sum of *t*-values exceeded 97.5 % (two-sided testing) of the permutation distribution. For the so defined carrier frequency cluster showing significant changes in amplitude coupling, we computed repeated measures ANOVAs with the within-subject factors condition (coherent vs. incoherent) and time (pre vs. post). To account for multiple comparisons, post-hoc *t*-tests were considered statistically significant if the corresponding *p*-value fell below the Bonferroni corrected alpha of *a_bonf_* = .025. To assess the stability of sustained coupling changes during the 5-minute post-stimulation period, repeated-measures correlations across subjects were computed on post minus pre AM-tACS amplitude coupling changes using 100-second segments with 50-second overlap after stimulation offset (Schwab et al., 2019).

The spatial pattern of sensor level AM-tACS effects was investigated by spatial cluster permutation statistics to correct for multiple comparisons. Clusters were obtained by the sum of *t*-values of electrodes which were adjacent in space and below an alpha level of 5 % (two-sided). The within condition permutation distributions were generated by randomly shuffling pre and post amplitude coupling values within participants, in each of 10.000 iterations. The between condition permutation distributions were generated by randomly shuffling amplitude coupling values between the incoherent and coherent condition within participants, in each of 10.000 iterations. The observed cluster was considered statistically significant when the sum of *t*-values exceeded 97.5 % of the permutation distribution.

To investigate whether observed amplitude coupling effects were paralleled by changes in power or phase coupling, the analysis steps described above for amplitude coupling were repeated for these parameters. The relation between amplitude coupling and power was further assessed using Pearson’s correlation coefficients.

In source space, we examined the relation between AM-tACS-induced electric field (E-field) strength and stimulation-related changes in amplitude coupling or power (post minus pre AM-tACS) using Pearson’s correlation coefficients. Analyses were confined to 177 cortical sources exhibiting E-field strengths ≥ 0.1 V/m, based on prior literature suggesting this threshold as sufficient to modulate neural activity and functional connectivity in humans (Jefferys et al., 2003; Kasten et al., 2019; Preisig & Hervais-Adelman, 2022).

To evaluate whether AM-tACS effects were biased by peripheral skin sensations or cognitive fatigue, both variables were transformed into ordinal Likert scales raging from 1 to 5. We analyzed the difference in peripheral sensations between AM-tACS conditions using Wilcoxon signed-rank tests. In addition, we examined the correlation between changes in amplitude coupling and corresponding changes in tactile sensation and cognitive fatigue using Kendall’s rank correlation coefficients.

## Results

### Incoherent AM-tACS decreases amplitude-coupling in the resting brain

To assess the effect of AM-tACS on interhemispheric amplitude coupling, we first computed the correlation of orthogonalized neural signals in the parieto-occipital electrode cluster, i.e., the target region of AM-tACS. The spectrum of amplitude coupling values across carrier frequencies showed a prominent peak in the alpha and low beta bands (Fig. 2A). This pattern replicates previous reports on the spectral structure of amplitude coupling during resting state in human electrophysiological recordings (Hipp et al., 2012; Siems et al., 2016; Siems & Siegel, 2020).

**Fig. 2.**
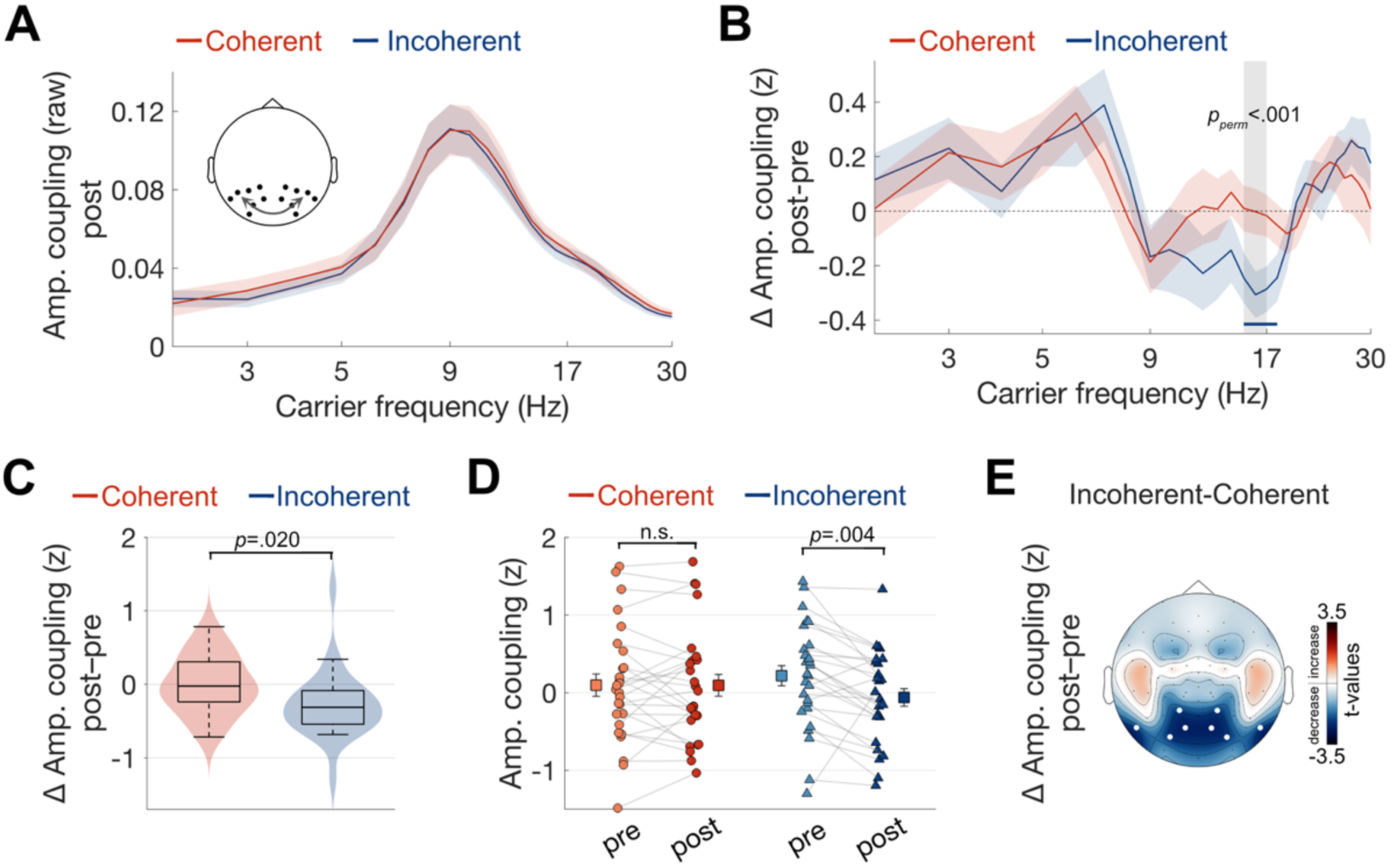
Interhemispheric amplitude coupling in the parieto-occipital cortex is significantly reduced by incoherent AM-tACS. (A) Spectrum of raw interhemispheric amplitude coupling averaged across post EEG resting-state recordings for the parieto-occipital electrode cluster. (B) Change in interhemispheric amplitude coupling (post minus pre AM-tACS) across carrier frequencies in the parieto-occipital target region. In a frequency cluster from 15-17 Hz, there was a statistically significant difference in amplitude coupling changes between AM-tACS conditions (grey shaded area). Additionally, there was a significant within-condition decrease in amplitude coupling by incoherent AM-tACS in a cluster from 15-18 Hz (blue line). (C) Box and violin plots illustrate the interaction effect within the 15-17 Hz frequency cluster, indicating a differential change in amplitude coupling (post minus pre AM-tACS) between AM-tACS conditions, with incoherent AM-tACS inducing a significant decrease. (D) Single-subject amplitude coupling values within the frequency cluster of 15-17 Hz pre and post AM-tACS. Squares indicate mean ± standard error of the mean. (E) Sensor level topography of changes in amplitude coupling between homologous electrode pairs within the 15-17 Hz frequency cluster. Statistically significant differences between AM-tACS-conditions were localized to an electrode cluster in the parieto-occipital cortex, i.e., the target region of AM-tACS.

To specifically test whether coherent and incoherent AM-tACS differentially modulated interhemispheric amplitude coupling, we examined changes in amplitude coupling across carrier frequencies for both AM-tACS conditions (Fig. 2B). Permutation statistics revealed a significant difference in the change of amplitude coupling between incoherent and coherent AM-tACS within a 15-17 Hz frequency cluster (*p_perm_* < .001). Furthermore, within-condition permutation tests showed that only incoherent AM-tACS induced a significant reduction in amplitude coupling within a cluster from 15-18 Hz (*p_perm_* < 0.001). Notably, this frequency range was centered on the carrier stimulation frequency of the AM-tACS signal of 17 Hz.

For the 15-17 Hz frequency cluster, a repeated measures ANOVA revealed a significant interaction between the factors condition and time (*F*(1, 26) = 6.19, *p* = .020, 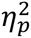 = 0.192; Fig. 2C), and a significant main effect of time (*F*(1, 26) = 5.41, *p* = .028, 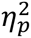 = 0.172). The main effect of condition did not reach significance (*F*(1, 26) = 0.02, *p* = .876, 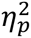 = 0.001). Post-hoc comparisons showed that the interaction effect was driven by a significant reduction in amplitude coupling following incoherent AM-tACS (*t*(26) = −3.20, *p* = .004, *d_z_* = −0.62), whereas coherent AM-tACS had no significant effect (*t*(26) = −0.04, *p* = .970, *d_z_* = −0.01) (Fig. 2D). Notably, the qualitative reduction in amplitude coupling under incoherent AM-tACS was constantly observed in 24 out of 27 participants, whereas changes under coherent stimulation were close to random, with 15 participants showing a reduction in amplitude coupling. To further evaluate the stability of stimulation-induced effects, we computed the changes in amplitude coupling within consecutive 100 sec epochs following AM-tACS offset. Repeated measures correlations across subjects revealed no significant change over time for either AM-tACS condition (coherent: *r* = .17, *p* = .079; incoherent: *r* = −.06, *p* = .537), indicating that AM-tACS effects on amplitude coupling persisted for at least the 5-minute post-stimulation period (see Supplementary Material A, Fig. A1).

As effects of electrical stimulation on neurophysiological activity were expected to be strongest in the AM-tACS target region, we assessed the spatial specificity of the observed amplitude coupling effect by spatial cluster permutation statistics. Fig. 2E shows the topographical distribution of condition-difference *t*-values for the change in amplitude coupling (post minus pre AM-tACS). Importantly, a significant difference between AM-tACS conditions in interhemispheric amplitude coupling changes was observed exclusively within an electrode cluster in the stimulated parieto-occipital target region (*p_perm_* = .020; for within-condition cluster permutations see Supplementary Material A, Fig. A2).

### AM-tACS effects are spatially specific to the parieto-occipital cortex and scale with E-field strength

To further assess the spatial specificity of AM-tACS effects, we used DICS linear beamforming to estimate source space activity and amplitude coupling between homologous cortical sources. As shown in Fig. 3A, over the entire cortex the reduction in interhemispheric amplitude coupling induced by incoherent AM-tACS was strongest within the parieto-occipital target region (for source level maps of power changes see Supplementary Material A, Fig. A3).

**Fig. 3.**
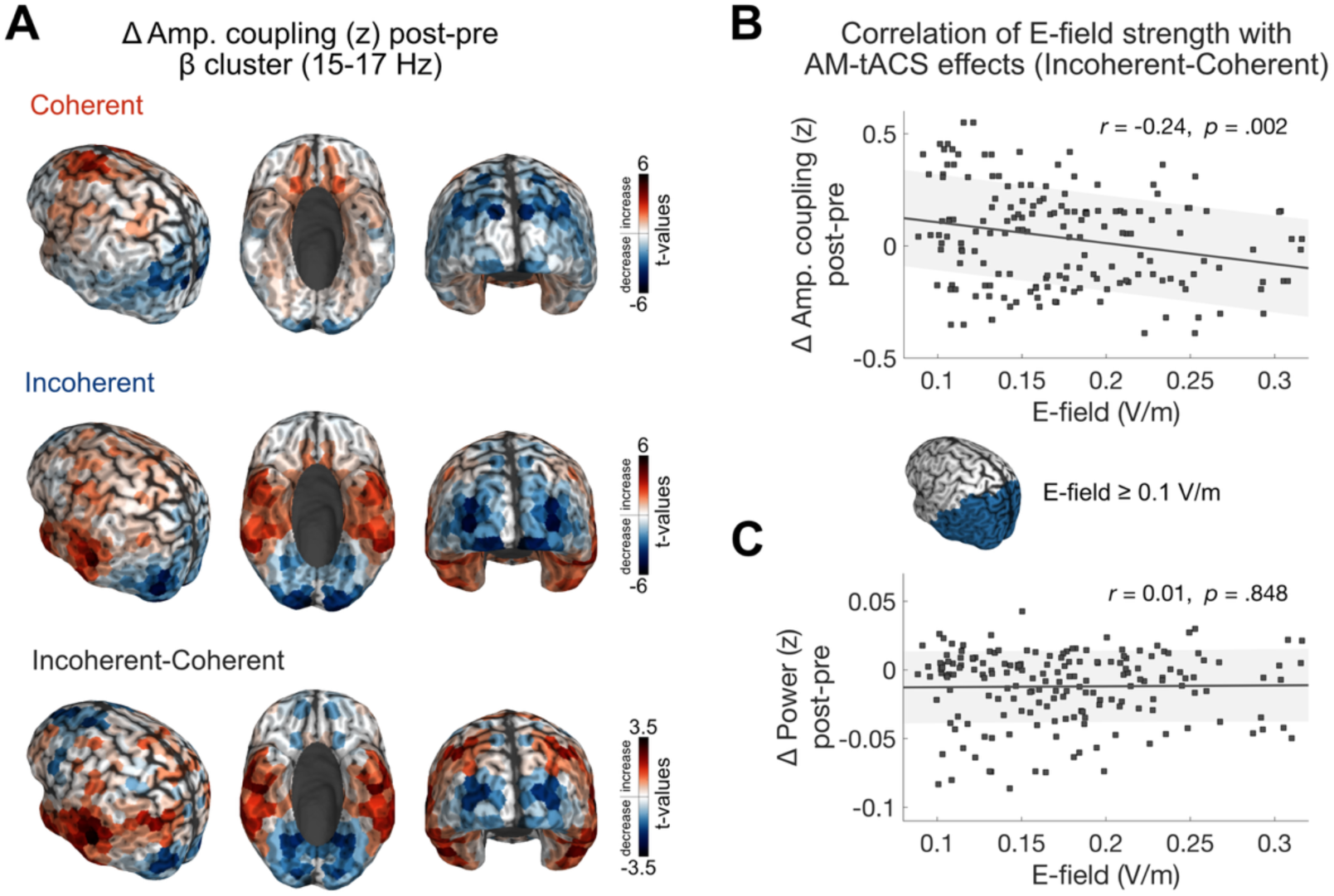
AM-tACS effects are spatially specific and linked to electric field strength. (A) Sources of AM-tACS-induced changes (post minus pre AM-tACS) in amplitude coupling shown for the coherent and incoherent condition as well as for their contrast (incoherent minus coherent). (B) The electric field (E-field) strength was negatively correlated with the differential change in amplitude coupling between incoherent and coherent AM-tACS within the frequency cluster from 15-17 Hz. This indicates that sources being exposed to stronger E-fields exhibit larger stimulation-dependent differences in coupling. (C) No significant correlation was observed between E-field strength and changes in oscillatory power. Correlation analysis was restricted to cortical sources exhibiting E-field strengths of 0.1 V/m or higher, a range in which neurophysiological effects of AM-tACS are expected.

If the bifocal electrical stimulation is directly linked to the identified after-effects on interhemispheric amplitude coupling, the effects should scale with the strength of the induced electric field. We first identified the cortical area exhibiting E-field strengths of 0.1 V/m or higher as a cortical cluster where neurophysiological effects of AM-tACS could be expected (Fig. 3B insert) (Jefferys et al., 2003; Kasten et al., 2019; Preisig & Hervais-Adelman, 2022). Notably, within this cortical cluster, AM-tACS-induced electric field strength was negatively correlated with the differential change in amplitude coupling between coherent and incoherent AM-tACS (*r* = −0.24, *p* = .002; Fig. 3B). Thus, cortical sources exposed to higher E-field magnitudes showed a stronger differential stimulation-induced change in amplitude coupling. This association was not observed for concurrent changes in oscillatory power (*r* = 0.01, *p* = .848; Fig. 3C), supporting the specificity of the AM-tACS stimulation effect on amplitude coupling, independent of local neural synchronization.

### Amplitude coupling effects are not mediated by concurrent changes in local oscillatory power

Changes in oscillatory power can influence the signal-to-noise ratio and, thereby, confound interpretations of amplitude coupling changes. We thus assessed the effects of AM-tACS on local power over the full electrode montage. In agreement with previously reported distributions of electrophysiological oscillatory power of the resting human brain (Capilla et al., 2022), occipito-parietal electrodes around the AM-tACS stimulation site displayed a prominent peak in alpha-band power (8–12 Hz). Further, the topography of beta-band activity (13–30 Hz, around the AM-tACS stimulation carrier frequency) showed a peak in occipito-parietal and sensorimotor areas (Fig. 4A and B). Thus, we next assessed whether AM-tACS decreased local power within the stimulation target region, which could have mediated the observed stimulation-induced reduction in amplitude coupling. Permutation statistics revealed no statistically significant difference in power changes between stimulation conditions. Within-condition cluster permutation statistics revealed a significant post-stimulation increase in power for coherent AM-tACS within the 4-8 Hz (*p_perm_* < .001) and 22-28 Hz (*p_perm_* < .001) frequency bands, and for incoherent AM-tACS within the 5-9 Hz (*p_perm_* < .001) frequency range (Fig. 4C). Importantly, these effects did not overlap with the beta frequency range in which amplitude coupling effects were most pronounced, i.e., 15-17 Hz. We further tested the power difference from pre- to post-stimulation within this frequency cluster using repeated measures ANOVA and did not find a main effect of stimulation condition or time nor an interaction effect (condition: F(1, 26) = 0.66, p = .423, 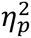 = 0.025; time: F(1, 26) = 0.03, p = .876, 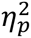 = 0.001; condition x time: *F*(1, 26) = 0.24, *p* = .630, 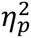 = 0.009) (Fig. 4D and E). In line with this result, spatial cluster permutation statistics did not reveal any significant electrode cluster for the differential change in power (post minus pre AM-tACS) between AM-tACS conditions (Fig. 4F; for within-condition cluster permutations see Supplementary Material A, Fig. A2).

**Fig. 4.**
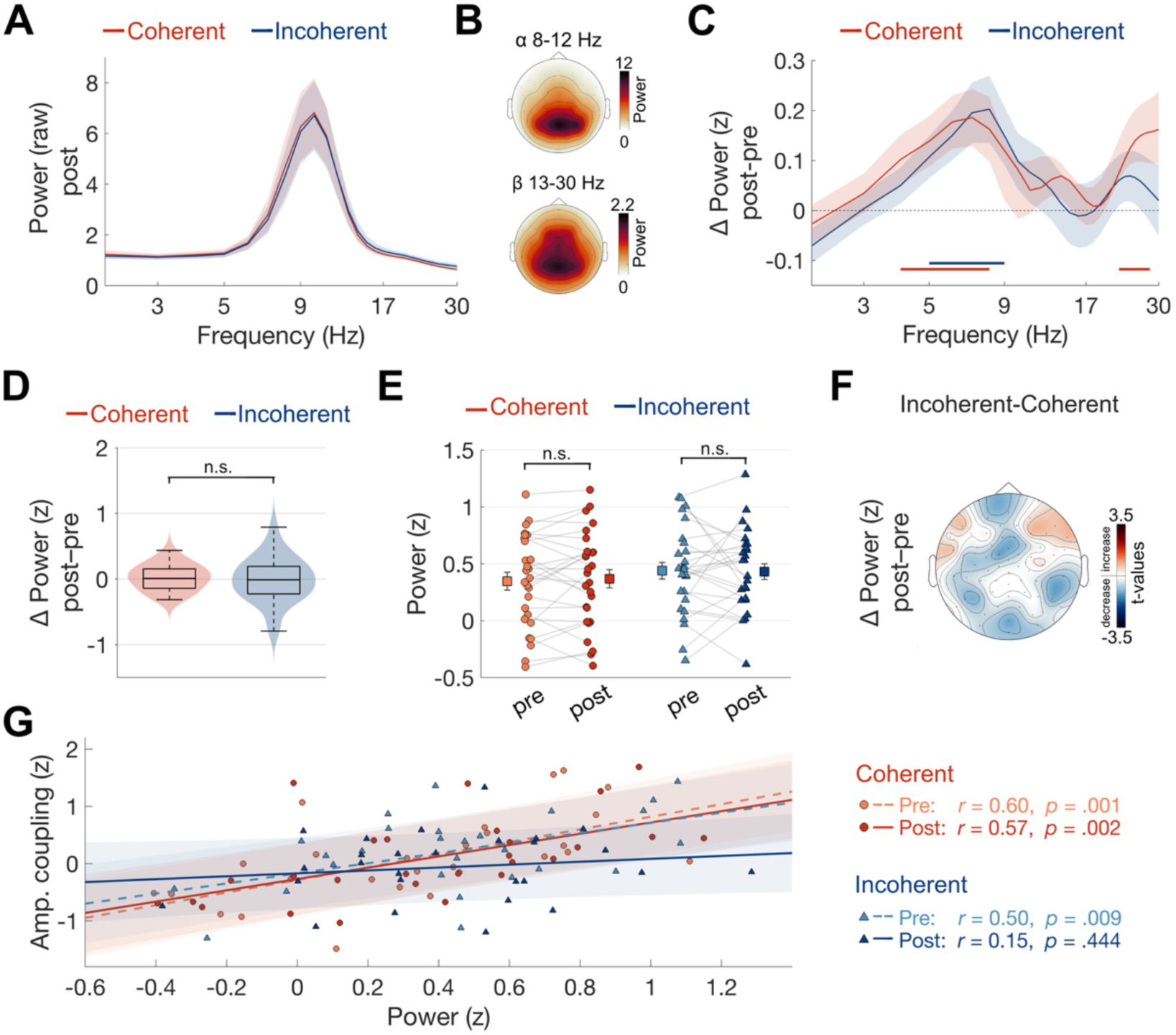
AM-tACS effects on amplitude coupling are not paralleled by changes in oscillatory beta power. (A) Power spectrum in the parieto-occipital target region across post EEG recordings for both AM-tACS conditions. (B) Topographies of alpha and beta power averaged across post EEG recordings for both AM-tACS conditions. (C) Change in power pre to post AM-tACS across carrier frequencies. The previously described AM-tACS effect on amplitude coupling in the beta frequency cluster from 15-17 Hz was not paralleled by changes in power, neither within, nor between AM-tACS conditions. Blue and purple lines mark frequency clusters showing significant within condition increases in power, i.e., in the theta and alpha range. (D) Box and violin plots illustrate the absence of an interaction effect, i.e., no difference in power change (post minus pre AM-tACS) between AM-tACS conditions within the 15-17 Hz frequency cluster. (E) Single subject data of power changes within the frequency cluster of 15-17 Hz. Squares indicate mean ± standard error of the mean. (F) Sensor level topography of the change in power within the 15-17 Hz frequency cluster. Permutation statistics reveal no significant effect. (G) Correlation between amplitude coupling and power for both AM-tACS conditions. The positive correlation structure between power and amplitude coupling is specifically disrupted post incoherent AM-tACS (dark blue, solid line). Shaded area indicates ±1 standard error of the estimated regression line.

Given that alterations in power are often associated with changes in amplitude coupling due to signal-to-noise-ratio changes, we directly analyzed the potential correlation between amplitude coupling and power for both AM-tACS conditions. As shown in Fig. 4G, a significant positive correlation was observed for the coherent AM-tACS condition, both before and after electrical stimulation (pre: *r* = .60, *p* = .001; post: *r* = .57, *p* = .002). In contrast, under incoherent AM-tACS, the positive pre-stimulation correlation was selectively disrupted following electrical stimulation (pre: *r* = .50, *p* = .009; post: *r* = .15, *p* = .444). Bayesian correlation test for the latter non-significant correlation coefficient yielded a Bayes factor of BF₁₀ = 0.25. This means that the observed disrupted correlation post incoherent AM-tACS is approximately 4 times more likely under the null hypothesis (no correlation) than under the alternative hypothesis (that the correlation exists). Together, these findings suggest that incoherent AM-tACS selectively disrupted interhemispheric amplitude coupling, while leaving local oscillatory power unaffected.

In addition to power analyses, we examined changes in phase coupling induced by AM-tACS. Consistent with the power results, no significant changes in phase coupling were observed within the beta frequency band, where amplitude coupling effects were most prominent (for details and statistics see Supplementary Material A, Fig. A4). Together, our findings indicate that incoherent AM-tACS disrupted interhemispheric amplitude coupling between the targeted cortical regions, independent of changes in local oscillatory activity. This highlights the capability of AM-tACS to selectively modulate amplitude coupling without affecting other forms of neural synchronization.

### AM-tACS effects on amplitude coupling cannot be explained by peripheral sensations

As electrical brain stimulation can activate cranial and cutaneous sensory nerves in the skin, thereby inducing tactile sensations in addition to direct transcranial effects, we applied EMLA cream to minimize peripheral sensory stimulation. Consequently, participants reported only mild to moderate skin sensations during AM-tACS with no significant differences in mean tactile sensation ratings (*W*(26) = 56.50, *p* = .115) or perceived cognitive fatigue (*W*(26) = 67.50, *p* = .089) between incoherent and coherent AM-tACS (Fig. 5A and B). If peripheral side effects of electrical stimulation accounted for the observed changes in amplitude coupling, one would expect the perceived intensity of tactile sensations or fatigue to correlate with neuromodulatory changes. However, no significant correlations were observed for either AM-tACS condition (all Kendall’s correlation coefficients *p* ≥ .225; Fig. 5C and D). Thus, although peripheral contribution to stimulation effects cannot be excluded, they were comparable between stimulation conditions and are unlikely to have driven observed effects on amplitude coupling, supporting a transcranial origin of AM-tACS effects on interhemispheric amplitude coupling.

**Fig. 5.**
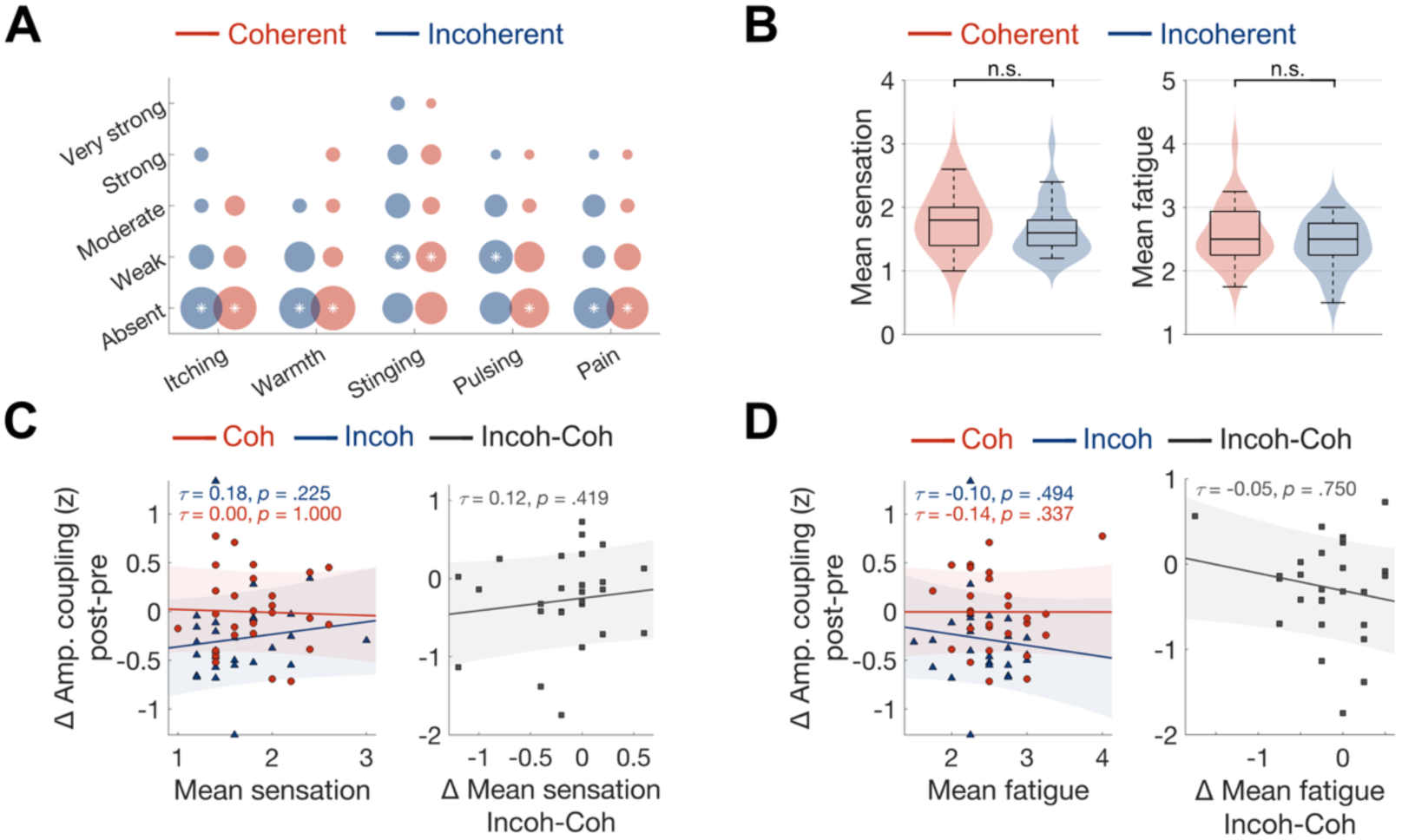
AM-tACS effects do not correlate with tactile sensations or cognitive fatigue. (A) Ratings of perceived intensity of tactile sensations (itching, warmth, stinging, pulsing, pain) for both AM-tACS conditions. The size of the circles indicates the overall count of ratings. Asterisks mark the median response. (B) Box and violin plots show the absence of condition differences in overall tactile sensation (left) or fatigue (right) ratings. (C) Left: The degree of amplitude coupling changes is not related to mean tactile sensation strength during coherent or incoherent AM-tACS. Right: No correlation between the difference between AM-tACS conditions in mean tactile sensation strength with the difference in amplitude coupling changes. (D) Left: There is no correlation between the degree of amplitude coupling changes and the level of cognitive fatigue for either AM-tACS condition. Right: No significant correlation between the difference between AM-tACS conditions in mean cognitive fatigue with the difference in amplitude coupling changes.

## Discussion

Our results demonstrate that dual-site AM-tACS can selectively and systematically modulate amplitude coupling during resting state in the human brain. By tuning AM-tACS envelope and carrier frequencies to physiologically plausible dynamics and applying AM-tACS with either coherent or incoherent envelopes between bilateral parieto-occipital cortices, we show that incoherent AM-tACS significantly reduces interhemispheric amplitude coupling for several minutes after stimulation offset. This effect was spatially confined to the stimulation target region and was most pronounced within the applied beta carrier frequency range. Furthermore, the amplitude coupling modulation strength was correlated with the induced electric field magnitude, suggesting a dose-response relationship with regard to the modulatory efficacy of AM-tACS. Importantly, changes in amplitude coupling occurred independently of alterations in local oscillatory power or phase coupling, emphasizing amplitude coupling as a distinct and dissociable mode of neural communication.

Intrinsic functional connectivity is commonly described by two principal coupling modes: amplitude coupling, reflecting slow co-modulations of oscillatory power, and phase coupling, capturing the fast synchronization of oscillatory cycles (Engel et al., 2001, 2013; Fries, 2015; Linkenkaer-Hansen et al., 2001; Palva & Palva, 2011; Siegel et al., 2012; Siems & Siegel, 2020; Singer, 1999). Whereas phase coupling has been linked to the transient coordination of neural processing, amplitude coupling is thought to index more stable, large-scale neural communication across distributed brain networks, thereby shaping and constraining faster, phase-based computations (Akam & Kullmann, 2012; Engel et al., 2013; Hipp et al., 2011; Lewis et al., 2016; Siegel et al., 2012). While the causal relevance of phase synchronization for perception and cognition has been established through a series of neurostimulation studies (e. g. Alekseichuk et al., 2017; Helfrich et al., 2014; Polanía et al., 2012; Reinhart, 2017; Reinhart & Nguyen, 2019; Strüber et al., 2014), proof for the causal role of amplitude coupling has remained scarce. By demonstrating that dual-site AM-tACS can selectively manipulate amplitude coupling independent of phase coupling, the present study addresses this gap and highlights the potential of AM-tACS as a flexible tool for experimentally dissecting the functional contribution of amplitude coupling to large-scale brain communication, cognition and behavior.

Crucially, observed modulations in amplitude coupling within the parieto-occipital target region were not accompanied by concurrent changes in oscillatory power that could confound the interpretation of amplitude coupling changes. Increases in power are typically associated with increases in amplitude coupling due to improved signal-to-noise ratios, a relationship reflected in the significant positive correlation between power and amplitude coupling observed before and after coherent AM-tACS, and well as before incoherent AM-tACS (Fig. 4G). Notably, this relationship was selectively disrupted following incoherent AM-tACS, indicating that incoherent stimulation modulated large-scale network communication independently of local oscillatory activity. Moreover, source level magnitude of the electric field is specifically predictive of the level of change in amplitude coupling, but not of changes in power. Taken together, these findings emphasize the specificity of AM-tACS for modulation of amplitude coupling, independent of other forms of neural synchronization, and support a dose-response relationship between the electric field strength and the modulatory potential of AM-tACS.

As concurrent EEG recordings during electrical stimulation are dominated by strong electrical artifacts, direct analyses of real-time stimulation effects are hampered. Therefore, we focused on the post-stimulation electrophysiological aftereffect of AM-tACS. While online effects may have been stronger, the outlasting effect of incoherent AM-tACS was sufficiently robust to be reliably detected in 24 out of 27 participants. The persistence of these effects, evidenced by stimulation-induced amplitude coupling changes that remained stable throughout the entire 5-minute post-stimulation period, points to lasting plastic adaptations within the targeted neural circuits, potentially mediated by spike-timing-dependent plasticity, long-term potentiation, or long-term depression (Kasten et al., 2016; Schwab et al., 2021; Vogeti et al., 2022; Vossen et al., 2015). Future work should examine longer post-stimulation intervals to further characterize the durability of AM-tACS-induced network changes.

The observation that incoherent AM-tACS significantly reduced amplitude coupling between the two AM-tACS target regions, whereas coherent stimulation had no measurable effect, reveals a differential responsiveness of neural circuits to these stimulation protocols. Converging evidence indicates that both the magnitude and direction of electrical stimulation effects critically depend on the individual brain state, with neural entrainment by tACS being most effective in individuals with low baseline activity (Alagapan et al., 2016; Krause et al., 2022; Neuling et al., 2013; Nguyen et al., 2018; Rufener et al., 2016; Ruhnau et al., 2016; Tseng et al., 2018). The simplistic dichotomy of coherent versus incoherent, or in-phase versus anti-phase stimulation, has further been challenged by a series of studies showing task- and brain state-dependent variability (Alekseichuk et al., 2017; Polanía et al., 2012; Reinhart, 2017; Tseng et al., 2018; Violante et al., 2017; Zhu et al., 2026). For example, Alekseichuk et al. (2017) reported that only anti-phase fronto-parietal theta tACS reduced phase coupling and impaired working memory performance, with in-phase tACS having no effect, whereas Schwab et al. (2019) found that in-phase bifocal occipito-parietal alpha-tACS increased phase coupling relative to anti-phase and jittered-phase stimulation. Further, Tseng and colleagues showed that in-phase theta tACS of bilateral parietal cortices improved working memory only in low performers, whereas anti-phase tACS deteriorated performance exclusively in high performers. Together, these findings suggest that the pre-existing strength of intrinsic oscillations and connectivity can constrain stimulation outcomes, potentially following an inverted U-shape relation. In our sample of healthy young participants, baseline amplitude coupling was likely optimally high (Hipp et al., 2012; Siems et al., 2016), limiting further enhancement by coherent AM-tACS due to ceiling effects. In contrast, incoherent AM-tACS was able to disrupt ongoing synchronization to induce a measurable reduction in amplitude coupling.

Building on this interpretation, our findings further support the framework that tACS may primarily reduce oscillatory power or coupling in healthy individuals with near-optimal neural dynamics as well as in disease-states marked by excessive neural synchrony, while neural enhancement may be more likely in individuals exhibiting suboptimal or dysregulated network states (Ahn et al., 2019; Lee et al., 2022; Marchesotti et al., 2020; Rufener et al., 2016). Accordingly, an important next step will be to assess the efficacy of incoherent versus coherent AM-tACS in patient populations with neurodegenerative disorders, such as Alzheimer’s or Parkinson’s disease, whose pathophysiology is characterized by reduced amplitude coupling dynamics and disrupted large-scale network organization (Boon et al., 2023; Schoonhoven et al., 2022; Stam et al., 2023; van Nifterick et al., 2024). Such work may help to clarify whether electrical brain stimulation can be used not only to perturb network activity but also to partially compensate for disease-related alterations and to restore amplitude-based connectivity.

As tACS has been shown to be most effective when the stimulation frequency aligns with endogenous neural rhythms (Fröhlich & McCormick, 2010; Ozen et al., 2010; Reato et al., 2010), we selected both the AM-tACS carrier and envelope frequency ranges to match envelope coupling dynamics typically observed during human resting-state recordings (Hipp et al., 2012; Siems et al., 2016). This physiologically informed approach was intended to maximize the physiological efficacy of AM-tACS in modulating ongoing amplitude coupling. The phase coherent beta-band carrier stimulation applied in both AM-tACS conditions may have stabilized ongoing beta oscillations, potentially providing a necessary foundation for modulating long-range amplitude coupling. Yet, whether the observed synchronization in amplitude co-fluctuations was indeed specific to the carrier signal frequency remains to be explored. Future work should systematically vary carrier frequency dynamics to clarify the frequency-specificity and reproducibility of AM-tACS effects.

In conclusion, our results provide the first electrophysiological evidence that dual-site AM-tACS can selectively and causally modulate interhemispheric amplitude coupling, independently of power and phase coupling, in a spatially specific manner. These findings establish AM-tACS as a powerful tool for manipulating amplitude coupling dynamics between distributed cortical regions, enabling the direct and causal assessment of amplitude coupling to cognition and behavior. Moreover, by supporting independent control over carrier and envelope components, AM-tACS provides a unique means to disentangle the respective roles of phase and amplitude coupling in large-scale brain communication and network organization. Together, our results highlight the potential of AM-tACS as a versatile approach for probing and modulating neural coupling, offering novel insights into how multiscale brain network dynamics support brain function.

## CRediT authorship contribution statement

**Marina Fiene:** Conceptualization, Methodology, Investigation, Formal analysis, Writing - original draft, Visualization, Data curation. **Marcus Siems:** Conceptualization, Methodology, Writing - review & editing. **Thorsten Kammerer:** Methodology, Investigation, Writing - review & editing. **Till R. Schneider:** Conceptualization, Methodology, Writing - review & editing, Supervision. **Andreas K. Engel:** Conceptualization, Methodology, Writing - review & editing, Funding acquisition, Project administration.

## Acknowledgements

We thank Karin Reimann and Viktoria Nickel for assistance with data collection, as well as all involved students for their participation.

## Supplementary Material A

**Fig. A1.**
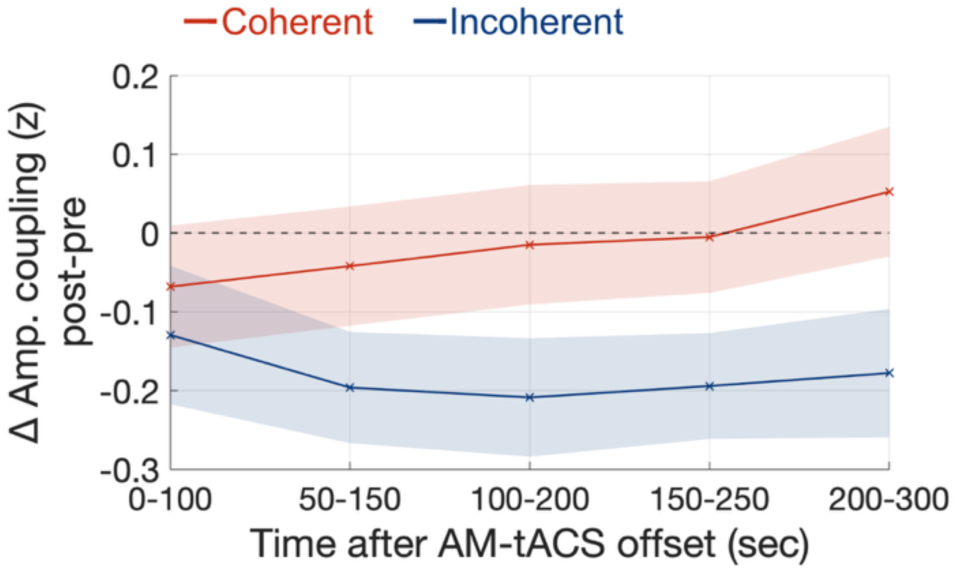
Stability of AM-tACS-induced aftereffects on amplitude coupling. Post minus pre AM-tACS changes in interhemispheric amplitude coupling within the 15-17 Hz frequency cluster for the coherent and incoherent stimulation condition. Repeated measures correlations across subjects revealed no significant change over time for either AM-tACS condition (coherent: *r* = .17, *p* = .079; incoherent: *r* = −.06, *p* = .537), indicating that stimulation-induced amplitude coupling changes remained stable throughout the entire 5-minute post-stimulation period.

**Fig. A2.**
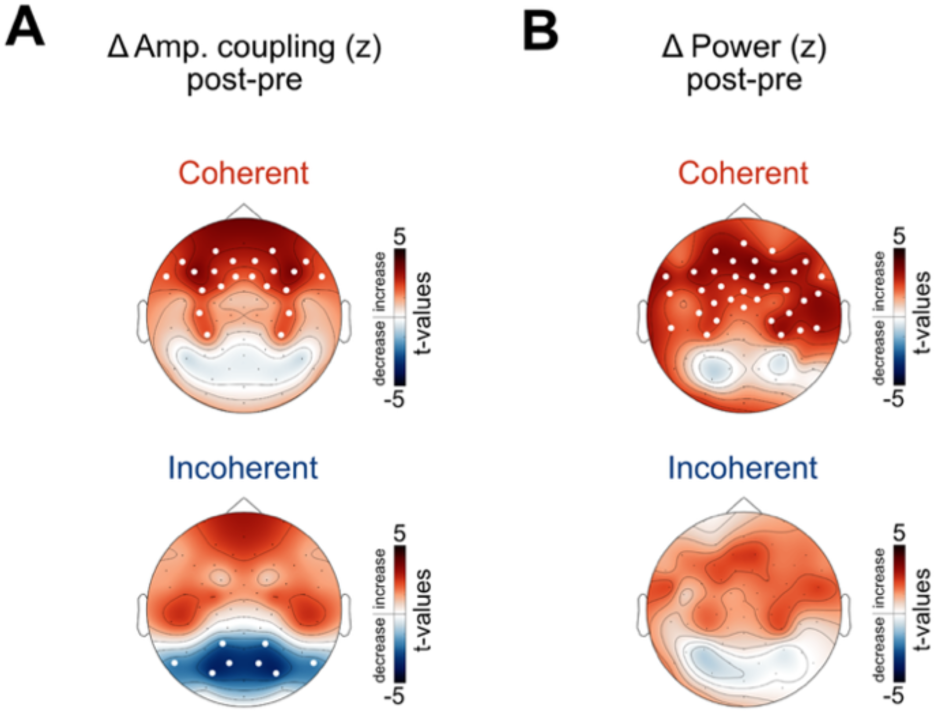
Sensor level AM-tACS effects on amplitude coupling and power for coherent and incoherent AM-tACS conditions. (A) Topography of post minus pre AM-tACS changes in amplitude coupling between homologous electrode pairs within the 15-17 Hz frequency cluster for the coherent and incoherent stimulation condition. (B) Topography of power changes within the 15-17 Hz frequency cluster for both AM-tACS conditions. For coherent AM-tACS, cluster permutation statistics revealed a significant increase in amplitude coupling in frontal brain areas (*p*_perm_ = .002), which was paralleled by a concurrent increase in power (*p*_perm_ = .008), precluding the interpretation of these amplitude coupling changes as genuine stimulation-induced effects on coupling. In contrast, for incoherent AM-tACS, statistically significant changes in amplitude coupling were localized exclusively to an electrode cluster in the parieto-occipital cortex (*p*_perm_ = .022), i.e., the target region of AM-tACS. This effect was not mediated by significant changes in power.

**Fig. A3.**
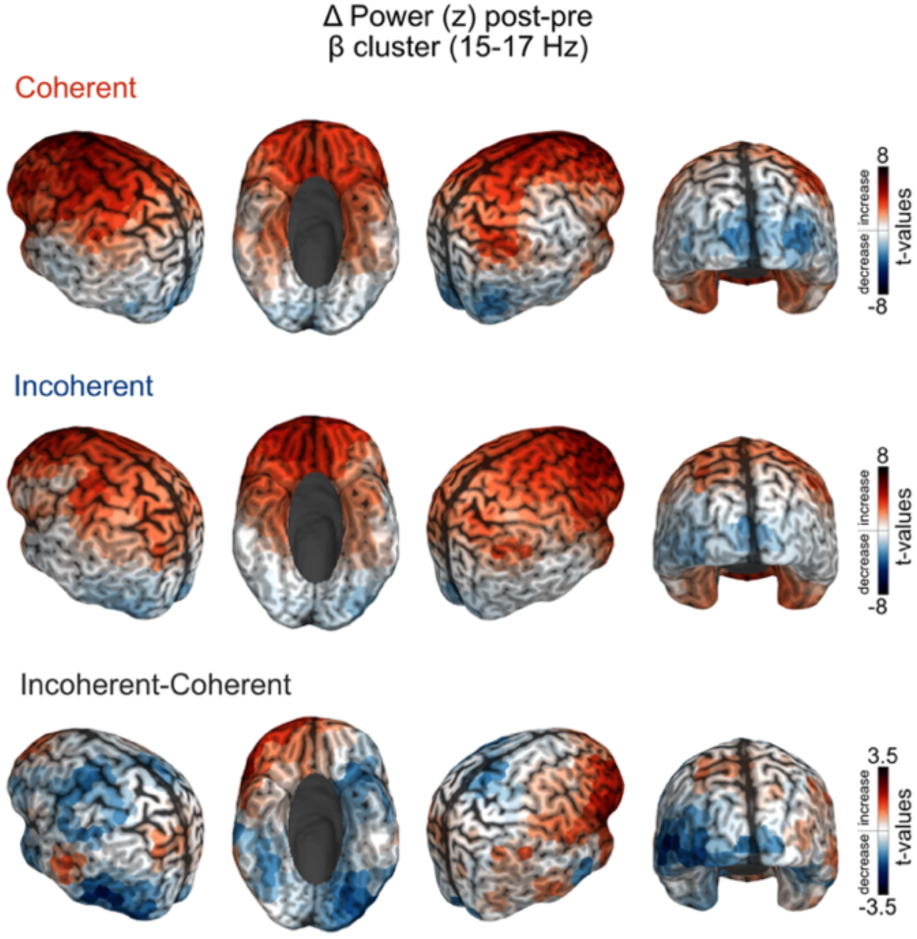
Source level AM-tACS effects on power. Sources of post minus pre AM-tACS changes in power within the 15-17 Hz frequency cluster shown for the coherent and incoherent AM-tACS condition, as well as for their contrast (incoherent minus coherent).

**Fig. A4.**
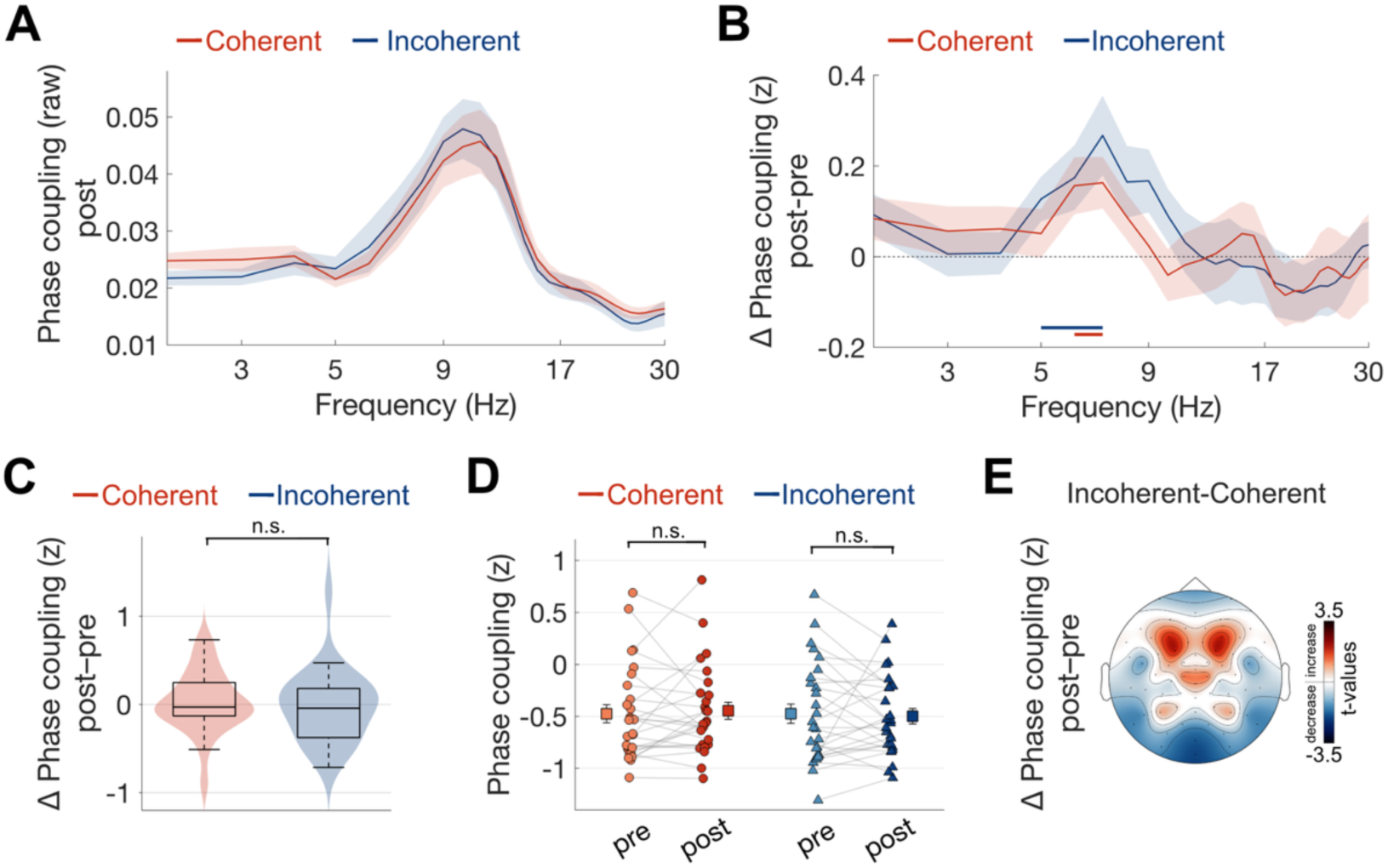
AM-tACS effects on phase coupling. (A) Spectrum of interhemispheric phase coupling across carrier frequencies in the parieto-occipital target region across post EEG recordings for both AM-tACS conditions. (B) Change in phase coupling pre to post AM-tACS across carrier frequencies. The AM-tACS effect on amplitude coupling in the 15-17 Hz frequency cluster was not paralleled by changes in phase coupling, neither within, nor between AM-tACS conditions. Blue and purple lines mark frequency clusters showing significant within condition increases in phase coupling, i.e., in the theta and alpha range (coh: 6-7 Hz frequency cluster, *p_perm_* = .009; incoh: 5-7 Hz frequency cluster, *p_perm_* < .001). Repeated measures ANOVA for the change in phase coupling within the 15-17 Hz frequency cluster revealed neither a significant main effect (condition: *F*(1, 26) = 0.14, *p* = .713, 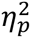 = 0.005; time: *F*(1, 26) = 0.001, *p* = .970, 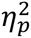 = 0.001) nor an interaction effect (*F*(1, 26) = 0.37, *p* = .547, 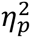 = 0.014). (C) Box and violin plots illustrate the absence of an interaction effect, i.e., no difference in phase coupling changes (post minus pre AM-tACS) between AM-tACS conditions within the 15-17 Hz frequency cluster. (D) Single subject data of phase coupling changes within the frequency cluster of 15-17 Hz. Squares indicate mean ± standard error of the mean. (E) Sensor level topography of the change in phase coupling within the frequency cluster of 15-17 Hz, with permutation statistics revealing no significant effects.

